# Short Tandem Repeat stutter_model_inferred from direct measurement of in vitro stutter noise

**DOI:** 10.1101/065110

**Authors:** Ofir Raz, Tamir Biezuner, Adam Spiro, Shiran Amir, Lilach Milo, Alon Titelman, Amos Onn, Noa Chapal-Ilani, Liming Tao, Tzipy Marx, Uriel Feige, Ehud Shapiro

## Abstract

Short tandem repeats (STRs) are polymorphic genomic loci valuable for various applications such as research, diagnostics and forensics. However, their polymorphic nature also introduces noise during *in vitro* amplification, making them difficult to analyze. Although it is possible to overcome stutter noise by using amplification-free library preparation, such protocols are presently incompatible with single cell analysis and with targeted-enrichment protocols. To address this challenge, we have designed a method for direct measurement of *in vitro* noise. Using a synthetic STR sequencing library, we have calibrated a Markov model for the prediction of stutter patterns at any amplification cycle. By employing this model, we have managed to genotype accurately cases of severe amplification bias, and biallelic STR signals, and validated our model for several high-fidelity PCR enzymes. Finally, we compared this model in the context of a naïve STR genotyping strategy against the state-of-the-art on a benchmark of single cells, demonstrating superior accuracy.

## Introduction

Short tandem repeats (STRs, also known as microsatellites) are repetitive elements of 1-6 base pairs long that constitute about 3% of the human genome. They are best known for their highly mutative properties *in vivo*, which is due to polymerase slippage that results in repeat contraction/expansion. Although their mutation rates vary dramatically, even low estimates are 3-4 orders of magnitude larger than of random point mutations, highlighting STRs as a tool of growing interest for various applications^1^. In disease, STRs are linked to tens of human diseases such as Huntington’s disease^2^; In several cancer types, mismatch repair deficiencies are analyzed utilizing STR polymorphic state, pointing to the disease progression^3^. In genetics studies, STRs are utilized to study population genetics and phylogenetics^4, 5^. In regulatory genomics, the importance of STRs as regulatory elements was recently demonstrated^6^. Recently, due to technological advancements in single cell (SC) genomics, SC STR analysis became of research interest for applications such as cell lineage phylogenetic analysis within an organism^7, 8^ and for pre-implantation genetic diagnosis^9^.

A key challenge for STR analysis is that they undergo noisy amplification *in vitro,* similarly to *in vivo* replication slippage. This noise, often termed “stutter”, is commonly manifested by excessive peaks when STR length data is plotted in a histogram of lengths (see example in **Error! Reference source not found.B**). Despite the value of the high polymorphicity of short unit STRs (e.g. in cancer diagnosis, forensics and phylogeny), they are still not commonly used for most assays due to excessive stutter noise. To address the stutter problem, simple noise models, such as highest peak analysis, are often employed. These simple models do not apply to highly polymorphic STRs, such as mono and di repeats, specifically in samples, which undergo substantial amplification. Using such models in these cases is likely to result in false genotyping. The problem of genotyping highly polymorphic STRs is even more difficult when genotyping non-hemizygous loci (such as from autosomal chromosomes, X Chromosome in female and in copy number variation (CNV) cases) since it is compounded by amplification imbalance of the two alleles. Such unbalanced amplification is typical in SC studies, as the starting material for WGA is a single copy of each locus.

With the growing need of *in vitro* amplification as a tool for basic and applicative scientific research, straightforward *in vitro* STR amplification studies were performed, in order to calibrate amplification factors and conditions ^5, 10–12^. A common STR stutter noise rule of thumb is that STR mutation rate both *in vivo* and *in vitro* is proportional to two main factors: (A) unit type length: short unit STRs (mono- and di-repeats) are more mutable than longer unit types. (B) STR length: Longer STRs (in repeat number) are more mutable than shorter STRs^1^. Nevertheless, despite years of STR research, a well-defined stutter behavior model is still lacking. The emergence of next generation sequencing (NGS) as a tool for large scale and detailed per-base analysis of STRs has re-emphasized the need for bioinformatics tools for STR analysis. While most current tools focus on mapping reads to the reference genome^5, 13, 14^, their stutter error correction algorithms are mainly calibrated with statistical models based on indirect measurements such as STR distributions in progenies, in populations and/or in user-defined data sets. Here we present a method for controlled measurements of stutter behavior during amplification for various STR types and sizes. Utilizing these measurements, we calibrated a mathematical model that accurately captures and predicts the stutter pattern of *in vitro* STR amplification.

## Results

### Controlled amplification of synthetic STR molecules

In order to study the stutter pattern as a function of amplification, we have designed and ordered a library of plasmids (**Error! Reference source not found.A**), each containing a unique combination of STR type and length, spanning all naturally occurring mono and di repeats (namely: A, C, AC, AG, AT) in the full spectrum of their natural genomic occurrence^15^ (Supplemental Table S1). The construct within each plasmid is sequencing-ready and includes a unique Illumina dual index combination for direct sequencing (T_1_) and a unique barcode for cross contamination control. Overall, the experimental setup allows for a controlled amplification and sequencing of all highly mutable STRs at three independent time points (T_1_-no amplification, T_2_-single amplification, T_3—_two amplifications) using various nested PCR primers, with the ability to measure the specific sequencing noise and bias for each STR length and type (**Error! Reference source not found**.B,C and Supplementary figures S1-S5).

### Fitting and model comparison

The data generated for the 3 time points (T_1_, T_2_, and T_3_) was used for the calibration of a computational model that predicts the stutter pattern at any theoretical amplification cycle given the repeat unit and length of the STR. Together with the assumption of perfect synthesis process (T_0_ – the designed construct prior to any manipulation), supported by Sanger sequencing.

Our goal is to predict the stutter histogram of repeat numbers for any amplification-time-point and for any original length in repeat units, and we assess the performance by:

Where is the distance between the two histograms. We have examined multiple distance metrics for the sake of histogram comparison and found 1-correlation^16^ to be the most suitable (Supplementary Figure S6).

We attempted to fit parameters to multiple models from the literature^17, 18^ and several in-house polynomial models by minimizing the overall distance between their resulting model histograms and the measured stutter patterns (Error! Reference source not found. B and Supplementary Figures S1-S5) using the Broyden–Fletcher–Goldfarb-Shanno (BFGS)^19^ optimization algorithm. We finally chose the model “Linear1up3dw”, a contraction-biased multistep linear model that best match the stepwise probabilities when implemented within a discrete-time Markov chain (Figure 2). This model obtained the best overall fit across the attempted mono and di STR when calibrated individually for each repeat type.

**Figure 1.**
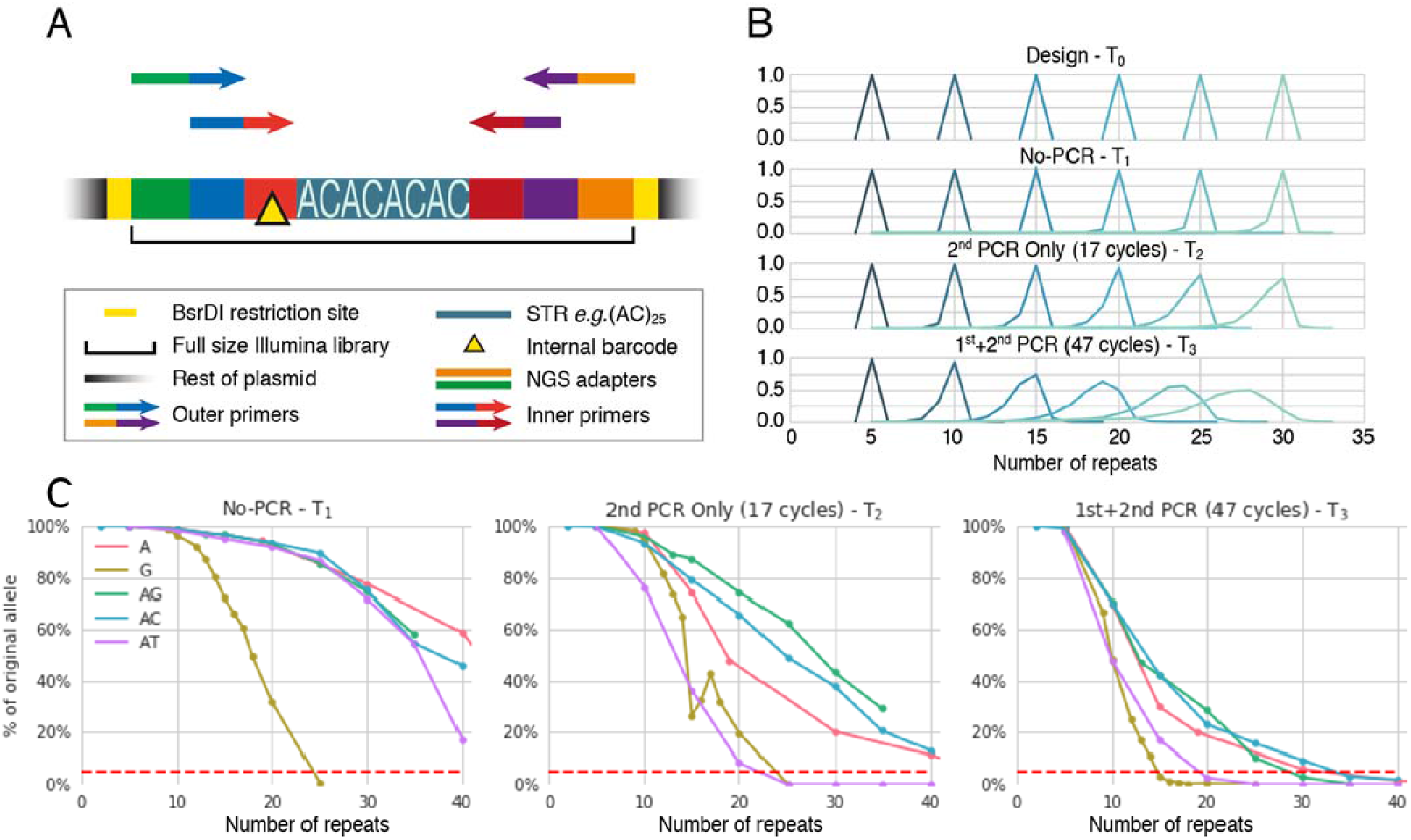
The synthetic STR experiment summary. **A**. Schematic description of the synthetic library. In each plasmid, a different synthetic STR construct was designed, synthesized and clone-sequenced for various STR types and length. The STR was designed within a context of an Illumina Truseq-HT dual index library to enable for nested PCR amplification at two time points (T_2_-amplification using outer primers only, T_3_-amplification using inner primers followed amplification by outer primers). The library is flanked by BsrDI restriction sites to enable direct sequencing of the STR library without amplification (T_1_). Internal barcode (yellow triangle) is a short sequence, unique to each STR length to detect for cross-contamination. See text and methods for elaboration and Supplemental Table S1 for the designed constructs. **B**. AC STRs repeat-number histograms, as were interpreted from sequencing results (T_1_, T_2_ and T_3_), compared to their expected length, T_0_ (designed sequence). **C**. Sequencing analysis results of each STR type, repeat-number and time point described as the percentage of the original (designed) signal from all the reads. Dashed line at the 5% marks the lower threshold of analysis: data points below the mark were deemed too noisy and were excluded from downstream analysis.

**Figure 2.**
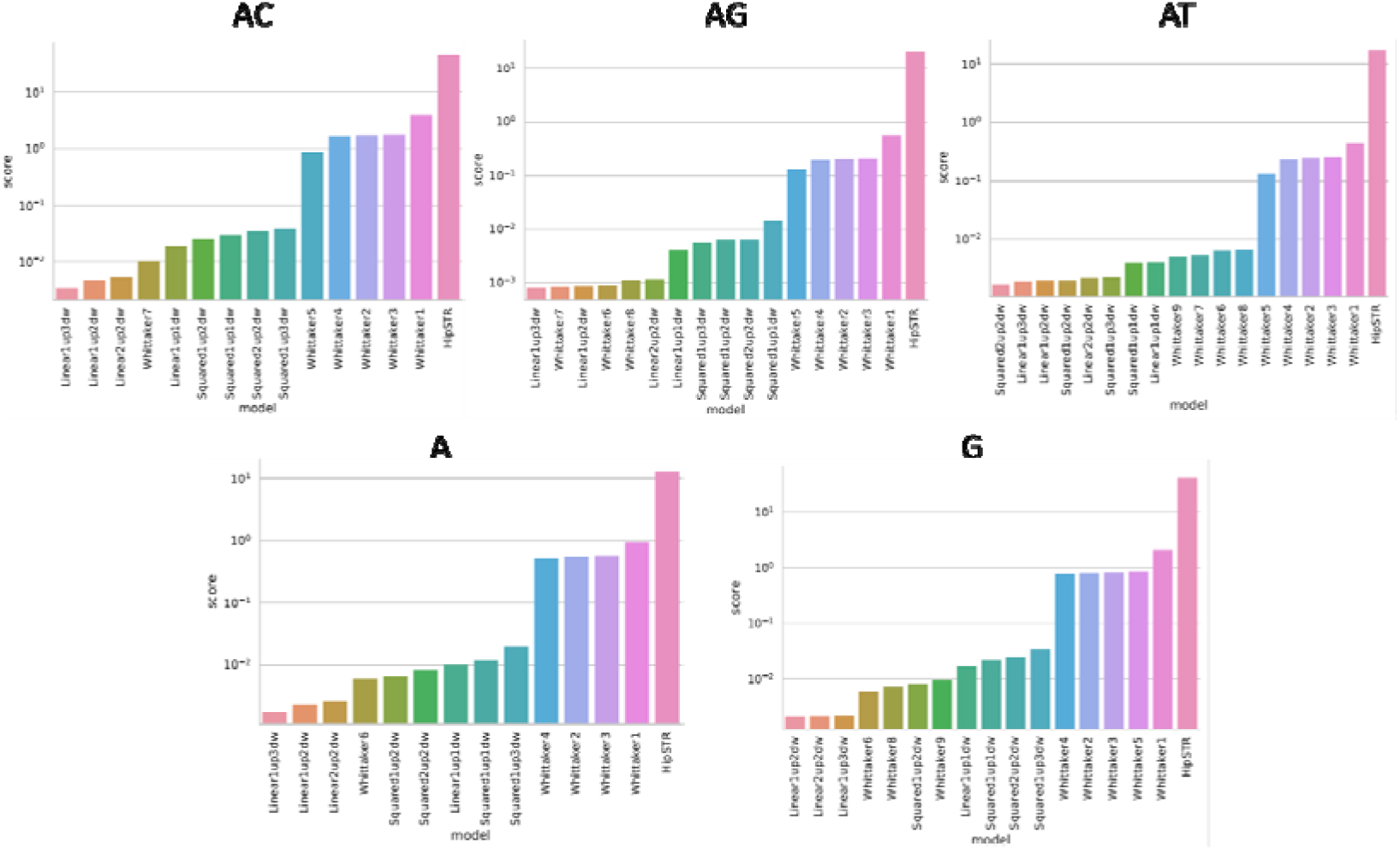
Model fitness to synthetic dataset. Each model parameters were optimized to best fit the dataset measured from the synthetic plasmids at different amplification time-points. The scores reflect the squared sum of 1-correlations across all measurements for each STR repeat unit type (*i.e.* AC, AG…) divided by the number of samples. Models include Whittaker1-9 as introduced by Whittaker *et al^17^*, HipSTR^18^ and polynomial models named after their number of variables and degrees.

We model these processes as an iterative mutation process with multiple steps. For each of these steps, our genotype can contract by up to 3 repeat units or elongate by a single repeat. The probability of such a mutation is linearly dependent on the STR’s current length.

### Validation and genotyping comparison

To confirm the model, we propose *R&B*, a naïve genotyping algorithm implementing an exhaustive strategy to call the original STR length from a population of reads with different STR lengths by scoring it against all possible predicted populations of any amplification time and STR length:

Following a meticulous STR genotyping comparison by Willems *et al.* ^18^, we compare this heuristic only to the current state-of-the-art, HipSTR genotyping tool, on a benchmark experiment first presented by Biezuner *et* al^8^. This experiment involves cells from a controlled *ex vivo* cell lineage tree experiment, picked and extracted for their DNA, while documenting their sampling lineage. STR mapping issues were tackled using an STR-targeted enrichment panel (rather than shotgun sequencing) and mapping the known primers panel to the reads in order to identify them. Using a similar strategy (FMSV^20^), we can isolate the problem of genotyping stutter patterns and avoid possible mapping bias.

The known lineage topology of individually analyzed SCs provides a solid reference for the comparison of any genotyping tool. To do so, we have devised the following metric to assess the accuracy of genotyping algorithms:

Let *A*: *T_leaves_* → *A* be the set of alleles assigned to the leaves of tree T by a genotyping algorithm. *P* (*A, T*) is the maximum parsimony or the minimal number of mutations required to explain set of alleles A on the leaves of tree *T*. 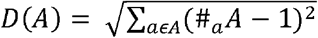 is the allele diversity.

We define F as the *reference tree fitting*.

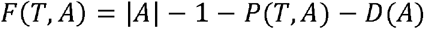

The *reference tree fitness* aims to balance the diversity of alleles found within this cell group, which provides information describing the topology of T, with the adherence of the genotypes to T. We compensate for the fact that diverse genotypes inherently have a lower parsimony, even when correct.

Using this metric, *Loci* that add valid information regarding the tree will be awarded positive scores while *loci* whose genotyping results contradict the topology will be negatively scored. A *locus* for which there is no relevant information (either no genotyping or a single allele across all cells) will receive a zero score.

Both genotyping methods, *R&B* and HipSTR, provide a measure of confidence together with each locus they attempt to genotype. While these confidence metrics are very different and have different distributions across the attempted cells/loci population, we can try to compare them by referring to percentiles of the full scores set, the top 10%, top 50% or any other threshold. To compare similar confidence genotyping attempts of both tools despite the large difference in the number of successfully genotyped loci, we compared only the cells/loci combinations for which both tools provide a genotyping attempt ranked with sufficient confidence (Figure 3A, B). Here we can see that across most confidence levels, when both tools attempt to provide a genotype, *R&B* attempts are more in line with the true tree topology.

**Figure 3.**
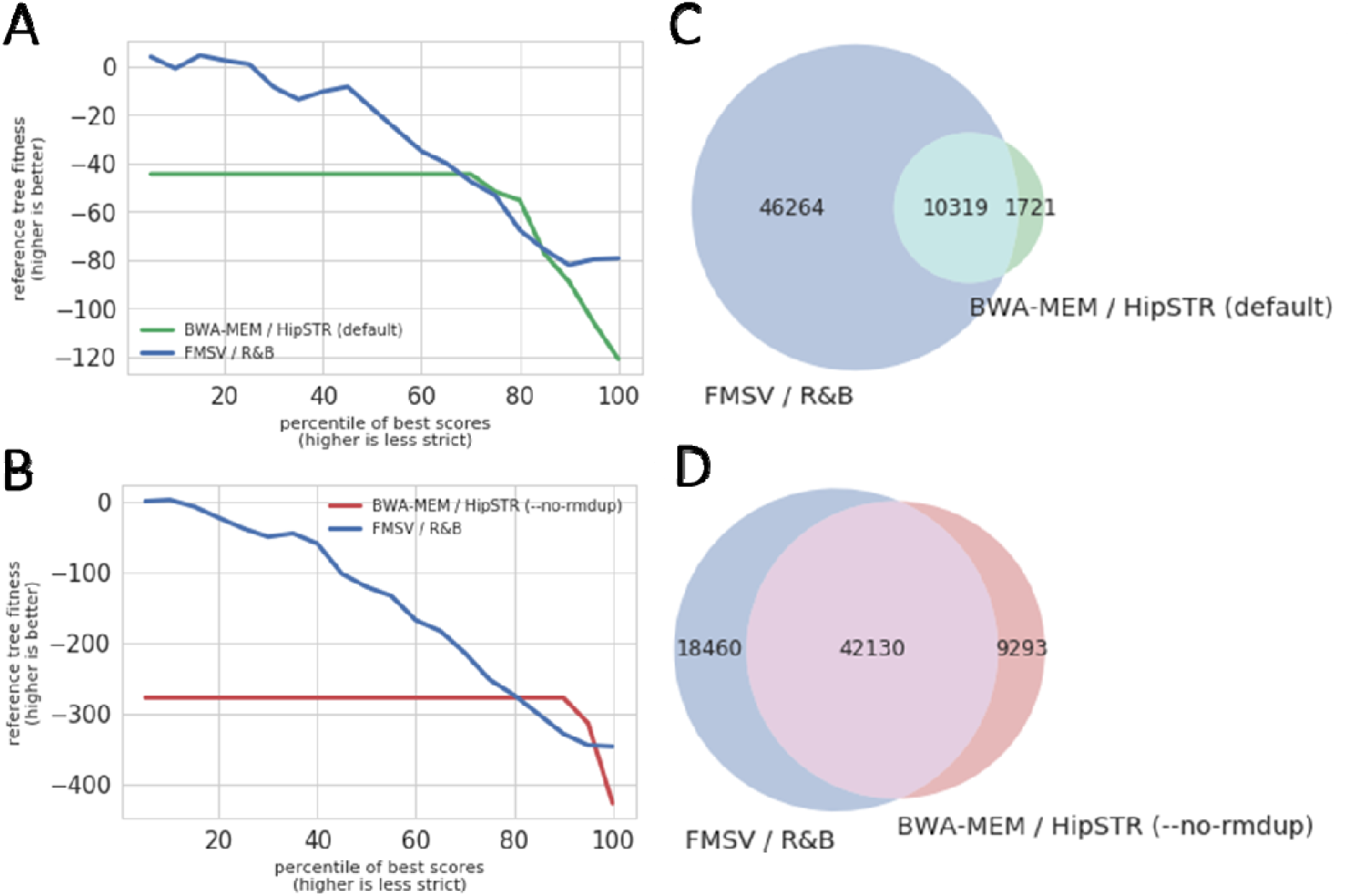
Genotyping results. Comparison of the proof of concept genotyping method, *R&B*, with HipSTR genotyping tool^18^ under default parameters and with the “--no-rmdup” flag, appropriate for PCR amplified results. We compare the results’ quality in tiles A and B by measuring their ability to accurately genotype the sequencing results of the *ex-vivo* cell lineage tree (reference tree fitness) as a function of their subjective confidence metrics (confidence greater than percentile threshold, lower values mean higher confidence but less loci). We compare loci that were genotyped by both genotyping methods within similar confidence percentiles (A, B) and the total quantity of the produced genotypes (C, D); We see that *R&B* excels in both quality and quantity. Across all cases, we used a minimal coverage of 5X, no confidence filters prior to percentile calculation and no stutter filtering for HipSTR. To maintain simplicity, we only account for haploid loci from the X chromosome of the cancerous cell line used in this experiment (human male DU145).

To maintain simplicity, we only account for mono-allelic loci from the X chromosome of the cancerous cell line used in this experiment (human male DU145). Other chromosomes were found to have major copy-number abnormalities.

## Experimental validation by controlled synthetic templates and real genomic data

To provide experimental-based confirmation for the model validity we opted to measure its simulated amplification cycles analysis in a series of controlled experiments. First, by using synthetic STRs as controlled templates for serial dilution analysis, and later, using synthetic STRs and real genomic data to demonstrate the robustness of the model analysis to the utilized PCR enzymes.

### Experimental validation of the model by using controlled synthetic templates

(1) Controlled amplifications of synthetic STRs in a serial dilution experiment. Using the synthetic STRs that were used for the model calibration above, we generated highly accurate NGS data originated from amplification of known and controlled templates. First, we have generated an NGS dataset generated from a single PCR amplification using the Q5 enzyme (NEB), as previously described for the T_2_ experiment, of 3 different templates: (AC)20, (AC)25 and (AC)30, each using 3 serially diluted templates (by 10 fold each). Our model’s simulated cycles linearly correlate with the actual number of amplification cycles performed, as expected from serially diluted samples (Supplementary Figure S7A, B).

(2) Model robustness to pcr enzyme by an enzyme comparison assay. First, we performed a small-scale PCR enzyme comparison by applying 5 commercially available PCR enzymes on the same synthetic templates as used above at an equal template concentration (using a subset of the generated data of Q5 from the abovementioned experiment (1) and 4 other enzymes). We show that the model accurately captures the stutter variability between different polymerases within a single degree of freedom, its simulated cycles (Supplementary Figure S7A, C).

### Experimental validation of the model by using single cell STR data

Following the successful proof of concept of polymerase comparison using synthetic templates, we opted to enlarge the validation to thousands of data points per each polymerase to create a statistical significant polymerase error rate comparative assay based upon the measured error rate per each thousands of genomic STR loci.

To generate a valid comparison we opted to utilize the same polymerase for the entire targeted amplicon sequencing amplicon protocol as outlined in ^8^ using 1769 amplicons (Supplemental Table S2). We first selected 6 high-fidelity enzymes and opted to apply them in parallel to a large collection of single cell WGA DNA template, on a large STR panel size in a 2-step amplicon targeted sequencing approach as described above. Specifically, the protocol follows a previously described method ^8^ which utilizes an AA chip for multiplex amplification and a 2^nd^ off-chip PCR for library barcoding, with the modification of PCR enzymes, thus, each sample will serve as template for all enzymes in a coupled 1^st^ and 2^nd^ PCR reactions. To fit all PCR enzymes in a single preliminary AA chip, we composed a “unified” 1^st^ PCR thermal cycler protocol that meets the requirements of all enzyme manuals (see methods section), with as little digression as possible from the manufacturers recommended protocols. 2^nd^ PCR was performed with each enzyme’s original protocol. The single cells taken from an H1 cell line which demonstrates a normal karyotype, thus reducing copy number artefacts (in this analysis, only X-chromosome STR were used). Following a preliminary analysis (Supplementary Table S3, Supplementary Figure S8) of two single cell DNA and 2 controls (positive and negative: bulk genomic DNA and DDW, respectively) in duplicates, we removed KOD enzyme from further analysis due to low success rate (mapped reads/total reads ratio). dNTpack, although failed in this experiment, since successfully utilized in the original protocol ^8^ with a different thermal cycler protocol was taken as a control for a large scale experiment. Here, we have enlarged the cohort of samples to 22 single cell DNA samples and 2 controls (positive and negative) were used as templates in an enzyme dedicated AA chip, in duplicates (48 samples per enzyme). Again, the 1^st^ PCR program was as the abovementioned “unified” and further PCR was in accordance with the enzyme protocol. dNTPack was used in two different AA chips: First (herein: “dNTPack”, as the rest of the enzyme (“unified” and than dedicated thermal cycler programs), an expected negative control. Second (herein: “UltraII/dNTPack”), as the 1^st^ PCR of the original AA protocol, in accordance with Fluidigm’s AA manual with cycle reduction as recommended by Fluidigm, and as was used in ^8^, with UltraII serving as its 2^nd^ PCR enzyme.

Applying our STR noise model on the sequencing data received from the large scale experiment we have generated an aggregated plot of simulated cycle scores of all single cells in the experiment (duplicates included, Figure 4A, see also results summary in Supplementary Table S4). We found that PrimeStar demonstrates a significantly lower simulated cycle number compared to the other enzymes. UltraII and Q5, both are second best in the number of simulated cycles category, demonstrating a similar result, thus emphasizing the robustness of the model, as both enzymes are based essentially on the same Q5 enzyme with a different mix composition. dNTPack shows the highest number of cycles amongst all examined enzymes. The original protocol (UltraII/dNTPack) demonstrates much higher success in number of successfully amplified loci count, making its 1^st^ PCR program preferable over the manufacturers protocol (in the context of utilization in AA chips), reasoning why the Fluidigm recommended such modification in the AA PCR protocol. Moreover, a cross between UltraII for the 2^nd^ PCR and dNTPack for the 1^st^ PCR showed improved score over dNTPack alone. These crossed results demonstrate the superiority of UltraII over dNTPack in noise insertion as could be expected by looking at the scores of each enzyme alone.

**Figure 4.**
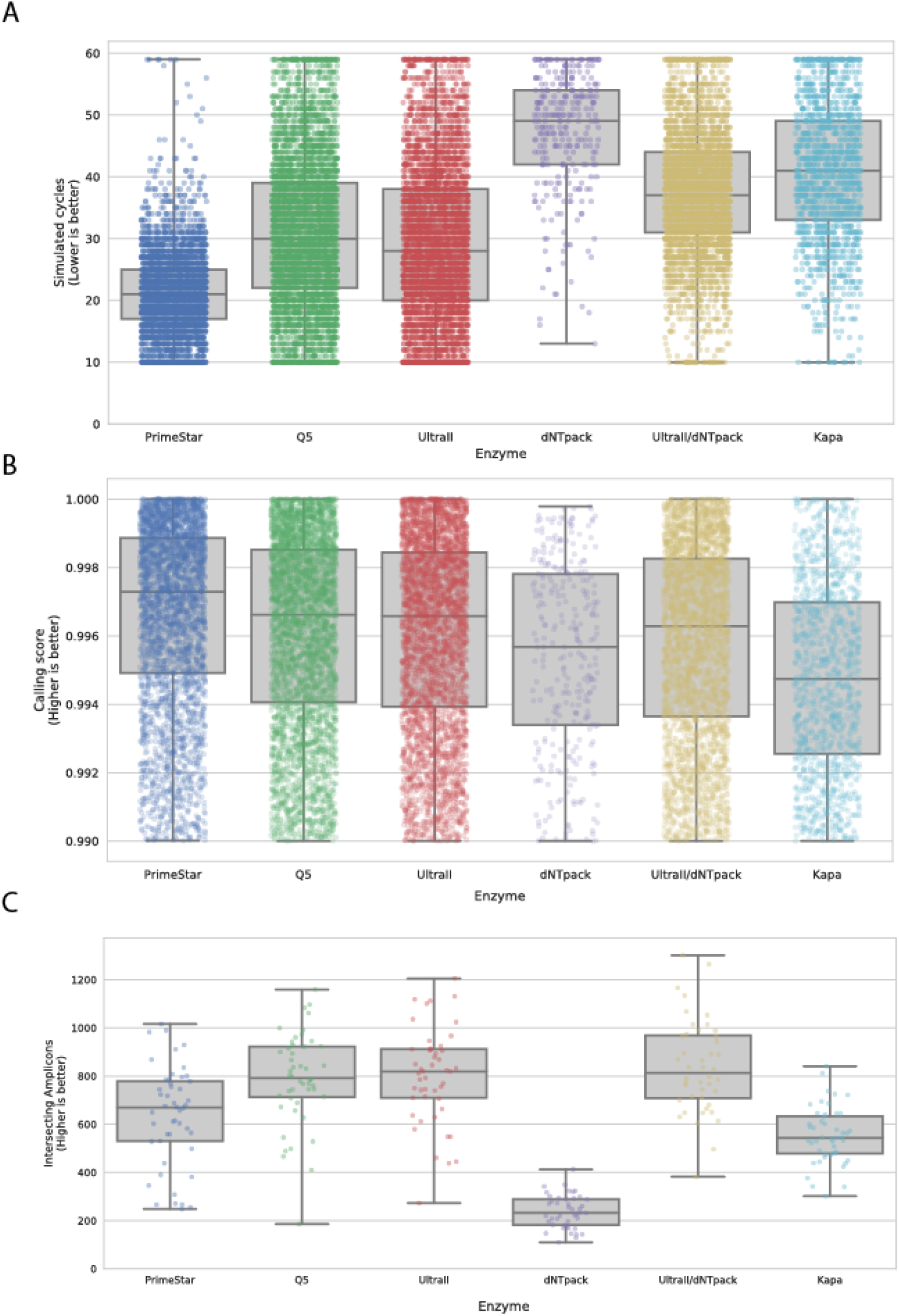
Comparison of genotyping results for various PCR enzymes using targeted PCR on a template of single cells WGA DNA. A) Comparing the number of simulated PCR cycles that best fit the measured histogram reflects the STR-specific stutter noise that is produced by a fixed number of actual PCR cycles. B) Comparing the fitness (correlation) between the simulated histograms and the measured ones. C) Loci counts that were retrieved from each SC.

Overall, we show that the model accurately captures the variability between different polymerases within a single degree of freedom, its simulated cycles, making it robust to any switch in utilized biochemical methods. It should be noted that although PrimeStar demonstrates the best score in noise insertion UltraII/dNTPack, as used in the current lab protocol (^8^, where Q5 is used instead of UltraII) shows good loci count(Figure XC), emphasizing that our current protocol works well in this category. However, UltraII and Q5 are also the best in the loci count category, and when considering both categories together, it seems that utilizing UltraII only, or PrimeStar only would be a preferable protocol, depends on the experiment requirements.

### Biallelic calling - genomic data

We opted to try and fit biallelic loci that amplified unevenly during the WGA process on SCs by extending the exhaustive search to nearly all possible allele combinations and at any proportion from the set: 0.1/0.9, 0.2/0.8,…,0.5/0.5…, 0.9/0.1 (Supplementary Figure S9). In order to assess our ability to accurately discover the true alleles that compose a stuttered biallelic histogram, we have selected autosomal loci from a SC population of H1 stem cells^8^ that consistently presented alleles A and B when genotyped as mono-allelic (Figure 5A,B,C first column). Since alleles A and B can appear at any proportion (Figure 5D), we can assume these cases presented the biallelic locus’ alleles at a proportion of 0/1 or 1/0 and that occurrences of this loci that failed to be genotyped as mono-allelic would present both alleles A and B.

**Figure 5.**
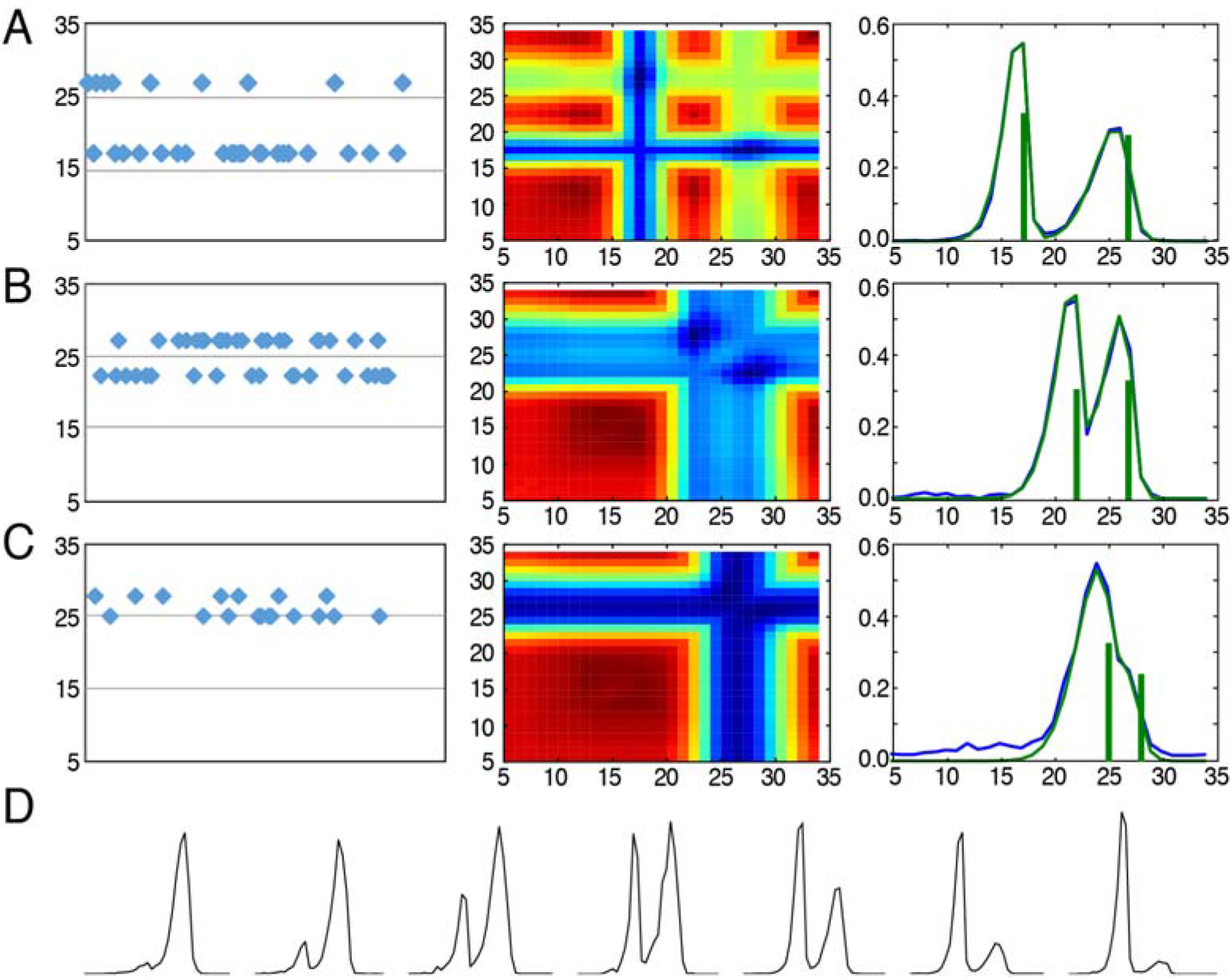
Biallelic genotyping using overlaid model histograms. Figure rows A, B and C show the successful genotyping of biallelic loci (AC repeats) within a SC population of H1 stem cells^8^. A, Recognizing overlapping alleles spanning 17 and 27 repeats, B, 22 and 27 repeats, and C, 25 and 28 repeats. First column – Monoallelic genotypes recognized in the clonal population. Second and third columns – In biallelic SC signal: Second column: Heatmap of the correlation scores between the predicted and the measured histograms across the space of possible alleles; Third column: Overlaid model prediction (green histogram) on top of the measured histogram (blue histogram). The resulting genotypes are marked as vertical green lines that also depict the alleles’ proportion in their height. D, Examples of asymmetric allele proportions.

## Discussion

STR usage in scientific research is increasing. High throughput sequencing opens a new frontier for STR science, both for basic^4, 6^ and for applicative research^21, 22^. With that understanding, in recent years, bioinformatics tools were developed to map and genotype STRs in a high-throughput genome-wide scale with improved accuracy and speed over standard mapping algorithms^5, 13, 14^. However, current tools still struggle with the *in vitro* amplification stutter noise that is typical to STRs, and in particular to highly mutable STRs. Recent biochemical advances have enabled PCR-free protocols that substantially decreased the effect of stutter noise in STR analysis^5^. However, these protocols have some limitations: (1) they require bulk amounts of template, making it incompatible with SC analysis, which requires whole genome amplification (2). In most cases, only a fraction of the STRs in the genome is required for analysis and therefore targeted amplification is required ^23^. Overall, this work lays the foundation for a better understanding of STR behavior in the NGS era. Although STR enrichment and sequencing kits are now available, a comprehensive assessment of the STR sequencing capabilities of extant sequencing machine was not systematically carried out, except for known constraints of some technologies such as mononucleotides sequencing in pyrosequencing based technologies^24^ and inferred estimation of such noise from old Illumina platforms^25^. Here we provided a controlled measurement of noisy sequencing at different amplification conditions and even in amplification free STR molecules.

We described here a new stutter model for the highly mutable STRs over *in vitro* amplification. The novelty of this model is that it is calibrated with NGS data generated by a controlled amplification of a range of di-repeat STRs of different types and sizes (according to their genomic occurrence in human). One key element in our model is that it takes into account that during amplification, the molecule lengths stochastic mutations can be accurately predicted, according to its inputs, the STR type, and the input length distribution of the previous amplification step. We chose to model the STR noise as a discrete-time Markov chain (DTMC). Our model enables easy calibration of different types of STRs. However, our data clearly shows a distinct and unusual pattern of noisy amplification of AT, which currently cannot be determined by either Markovian or binomial models, and may require modified model in the future. This variation in mutational mechanism was suggested previously^1^.

We provided three types of experimental-based evidence for the effectiveness of our model:

(1) Controlled amplification of STR plasmids. First, by utilizing it to measure an accurate amplification difference between known STR templates of various types and concentration, and second, by validating it against various types of polymerases.
(2) Comparative analysis of STR amplification of thousands of genomic single cell STRs. Both experiments have demonstrated the model robustness, such that although calibrated by a specific set of polymerases and conditions can be trustfully used as a quantitative tool for analyzing mutational processes by any NGS downstream process. Future work will enable a large-scale utilization of this model for assaying and/or optimizing other mutational processes, such as WGA.
(3) Utilization of NGS genomics datasets from SCs by accurately analyzing STRs from biallelic histograms, from drifted histogram, unclear determination of single peaks, and unbalanced allelic representation.

We also compared our model to a state-of-the-art genotyping tool^14^ utilizing NGS data from SC targeted enrichment data, originated from an *ex vivo* controlled cell lineage tree^8^. Our model outperforms both by the number of STR genotypes and both by the calling confidence, when compared with respect to the *ex vivo* tree.

We acknowledge that the bioinformatic improvement we provide here is mainly the stutter model itself, where current tools, mainly HipSTR, are implemented as a more inclusive STR genotyping tools in terms of phasing, haplotyping and interfaces with standard bioinformatics pipelines. Nevertheless, we recommend this model as an integrative step for STR noise analysis, specifically for SC analysis, where the sequenced samples undergo extensive amplification or in high sensitivity STR analysis, e.g. diagnosis of Microsatellite Instability (MSI) in cancer samples^26^. The tolerance of our model to noisy STR signal allows for a more flexible experimental design and opens the gate for highly mutable STR sequencing research.

## Materials and Methods

### Controlled amplification noise measurement of a synthetic STR library

STR plasmid design: Sequence verified cloned plasmids containing synthetic STRs of different types and sizes (Supplemental Table S1) were ordered from either IDT or GenScript (pIDT-kan and modified puc57-Kan vectors, respectively). Cloning vectors were validated to exclude BsrDI restriction sites. STRs were synthesized in the context of a complete Illumina NGS library (Truseq HT) to allow for nested amplification, and to enable a direct digestion using the Type IIS restriction enzyme BsrDI, thus creating a sequencing ready library. See elaboration in main text and in **Error! Reference source not found**. Immediate STR flanking sequences were validated to avoid partial STR repeat unit occurrence *(e.g.* (AC)_X_ followed by “A”). Internal 3-mer internal barcodes were inserted to allow for cross-contamination detection between samples. Several amplification time points were measured:

#### Ti (No-PCR) control

was performed by pooling all STR plasmid libraries at equal concentration and digestion with BsrDI enzyme (NEB) according to manufacturer protocol. Digestion was performed at 65°C for 16 hours, followed by inactivation at 80°C for 20 minutes. Reaction was then processed for sequencing (see later description in “Pooling and sequencing”).

#### T_2_ and T_3_ PCR experiments

In the T_3_ experiment, each STR plasmid (10^−4^ μg/μl) was loaded as template in an AccessArray (AA) PCR chip. Each primer inlet was loaded with the same primer solution (“Inner primers”) composed of X1 Access Array Loading Reagent (Fluidigm) and primers: Control_Fw: 5’-CTACACGACGCTCTTCCGATCTTCCTAATCTTACGCGGCCATAAC-3 ‘ and Control_Rev: 5’-CAGACGTGTGCTCTTCCGATCATGGACAGTCTTTAAGAGCCCATC-3’(IDT), at a concentration of 1μM each. PCR reactions and purifications were performed as described in^8^: In summary, a 1^st^ PCR of 30 cycles PCR reaction is performed in the AA chip. Following sample harvesting, purification and dilution 1:100, a 2-step 2^nd^ PCR of 17 cycles (5 cycles with annealing temperature of 55°C + 12 cycles with annealing temperature of 70°C) is performed to generate a dual indexed sequencing library (note that the “Outer primers” sequences were as described for the 2^nd^ PCR primer sequences in^8^. The 1^st^ PCR (in the AA chip) is done using the manufacture recommended enzyme: FastStart High Fidelity PCR System, dNTPack (Roche) while the 2^nd^ PCR is done using Q5 Hot Start High-Fidelity DNA Polymerase (NEB) with the addition of SYBR green I (LONZA) at a final concentration of X1, to enable real time tracking of amplification. Following 2^nd^ PCR, each sample was purified using SPRI beads. T_2_ PCR was performed by using 0.1ng-1ng of each STR plasmid as a template. Samples were processed in accordance with the T_3_ 2^nd^ PCR protocol.

#### Pooling and sequencing

All samples (T_1_, T_2_, T_3_) were purified and concentrated using MinElute PCR purification kit (Qiagen), pooled together and size selected (200-500bp) using a 2% agarose BluePippin gel cassette (Sage Science) utilizing an upgraded software that avoids blue light exposure after marker detection. Products were concentrated again (Minelute) and were sequenced by a 2×220bp sequencing (Miseq, Illumina) using a manufacture recommended sequencing primers (R1, Index) and custom R2 primer 5’ -GTGACTGGAGTTCAGACGTGTGCTCTTCCGATC-3’ (HPLC grade, IDT).

### Experimental validation of the model by using controlled synthetic templates

We opted to validate the model using five high fidelity PCR enzymes, using the controlled synthetic STRs as templates. The enzymes were: the two enzymes that were described above (Q5 High-Fidelity DNA Polymerase and FastStart High Fidelity PCR System, dNTPack), Phusion High-Fidelity DNA Polymerase (NEB), KOD Hot Start DNA Polymerase (Novagen) and KAPA HiFi HotStart PCR Kit (Kapa Biosystems).

Reactions were as performed in the T_2_, described above: 20μl reactions in a 96-well format, with real time amplification tracking using SYBR green I, each time using a different enzyme and buffer composition, different templates, and different barcoding primers. The template for each PCR was 2μl of 1ng/μl STR plasmids: (AC)_20_, (AC)_25_, or (AC)30. Each reaction was duplicated to avoid PCR primer sequence effect (using different indexes). Negative control (water) was added to each PCR. In the serial dilution validation experiment, Q5 enzyme was used as described above, using the same STR plasmids as templates in 3 concentrations: 1 ng/μl (also used for the enzyme comparison experiment), 10^−2^ ng/μl and 10^−4^ ng/μl.

All Samples were purified, pooled and sequenced as described above. The following exceptions were considered: 1) Activation, elongation and final elongation were adjusted to fit each enzyme’s recommended protocol. 2) Annealing temperature from the 6th amplification step and on was according to each enzyme’s elongation temperature. 3) PCR reaction was stopped when amplification reached a plateau. 4) Due to failure of dNTPack to amplify using the standard 2-step PCR protocol, we applied the same program as being performed in the 1^st^ PCR of T_3_ (in the AA chip). 5) Reactions mixes were according to manufacturer’s protocols, with primer concentrations of 0.3-0.5μM, with the exception of dNTPack, which composition was according to Fluidigm’s recommended reaction mixture with primer concentration of 0.1μM each and a final volume of 10.6μl.

### Experimental validation of the model by using single cell STR data

The high fidelity PCR enzymes were used in this study were: NEBNext Q5 Hot Start HiFi PCR Master Mix (NEB), NEBNext Ultra II Q5 Master Mix (NEB), FastStart High Fidelity PCR System, dNTPack (Roche), KOD Hot Start DNA Polymerase (Novagen), KAPA HiFi HotStart PCR Kit (Kapa Biosystems) and PrimeStar Max (Takara).

A recreation of the original amplicon targeted sequencing protocol as presented in ^8^ was performed in order to assess the error rate per polymerase enzyme using the STR stutter model. In summary, AA chip generates a mixture of 48 X sample+PCR wells, with 48 X primer mixes (1769 of amplicons in total, see Supplementary Table S2), ending up with 2304 nanoliter reactions, which are later harvested to each sample’s inlets (48 reactions to a single well). Following sample harvesting, purification and dilution 1:100, a 2^nd^ PCR was performed at a final volume of 20μl, each sample with its corresponding PCR enzyme from the 1^st^ PCR reaction and using its protocol, unless otherwise mentioned. Purification and pooling procedures were as described in^8^. The “unified” AA 1^st^ PCR protocol as abovementioned described was composed in accordance with the thermal cycling protocol guidelines of all examined polymerases: Activation was performed at 98°C for 3 minutes followed by 5 cycles of 98°C for 20 seconds, 60°C for 15 seconds and 70°C for 15 seconds, and 20 cycles of 98°C for 20 seconds, 70°C for 15 seconds and 70°C for 15 seconds. A final elongation step was added: 70°C for 5 minutes. Each polymerase reaction mixture was according to its manual. To avoid over-cycling SYBR green (X1) was added to enable amplification tracking by real time PCR. Libraries were first shallow sequenced in 2 × 220-bp in a Miseq sequencer (Illumina), followed by normalization and pooling by number of total reads per sample, and deep-sequenced in NextSeq(Illumina) 2 × 151-bp.

### Computational analysis

For the initial analysis of the mini-genes library, enzyme comparison and biallelic genotyping, the pipeline presented by Biezuner *et al*^8^ was used. In short, reads are processed using *cutadapt* (https://cutadapt.readthedocs.io/en/stable/) and PEAR (<u>Zhang et al. 2014</u>), followed by unique mapping of the merged reads to their target using read alignment of only the read’s edges corresponding to the primer pairs. STR repeat number is then determined by aligning the read to references containing a range of STR lengths and choosing the reference length with the highest alignment score.

For the initial analysis of the library that was used for comparing genotyping accuracy, FMSV^27^ mapping was used to generate the input for *R&B* genotyping tool while BWA-MEM was used to generate the input for HipSTR^18^.

## Acknowledgements

We thank O. Bechar for the prompt design of figures. This research was supported by The European Union grants: ERC-2008-AdG (Project No: 233047) and ERC-2014-AdG (Project No: 670535); by The Israel Science Foundation grants: Individual Research Grant (Grant No: 456/13) and Joint Broad-ISF Research Grants: 422/14 and 2012/15; by The German Research Foundation DFG SH grant 867/1-1 and by The Kenneth and Sally Leafman Appelbaum Discovery Fund. E.S. is the Incumbent of The Harry Weinrebe Professorial Chair of Computer Science and Biology.

